# Prediction and final temporal errors are used for trial-to-trial motor corrections

**DOI:** 10.1101/368001

**Authors:** Joan López-Moliner, Cécile Vullings, Laurent Madelain, Robert J. van Beers

## Abstract

Many daily life situations (e.g. dodging an approaching object or hitting a moving target) require people to correct planning of future movements based on previous temporal errors. However, the actual temporal error can be difficult to perceive: imagine a baseball batter that swings and misses a fastball. Here we show that in such situations people can use an internal error signal to make corrections in the next trial. This signal is based on the discrepancy between the actual and the planned action onset time: the prediction error. In this study, we used three interception tasks: reaching movements, saccadic eye movements and a button press that released a cursor moving ballistically for a fixed time. We found that action onset depended on the previous temporal error in the arm movement experiment only and not in the saccadic and button press experiments. However, this dependency was modulated by the movement time: faster arm movements depended less on the previous actual temporal error. An analysis using a Kalman filter confirmed that people used the prediction error rather than the previous temporal error for trial-by-trial corrections in fast arm movements, saccades and button press.

## Introduction

Timing errors of actions are ubiquitous in daily life and learning from these errors to improve planning of future movements is of great importance. Suppose you are batting in a baseball game and you just missed a fastball by 50 ms. Assuming you validly expect another fastball, how and how much should you correct for this error in the next movement may depend on different factors. You could use an estimate of this temporal error (between the bat and the ball) and try to react earlier if you were late but your perception of this error will be noisy. Since the movement time of your hitting movement can be quite constant you could alternatively rely on correcting the onset of the swing relative to some relevant moment (e.g. ball motion onset). In this study, we ask on what basis one corrects for temporal errors under various situations of uncertainty regarding the final temporal error and the possibility of correction during the movement. Correcting on the basis of previous errors is one of the hallmarks of motor learning (1,2) and many studies have addressed how people correct for spatial errors in response to some external perturbation (e.g. with force-fields or distorted visual feedback) (3-7) or in situations without perturbations (8).

It is known that larger uncertainty on the perceived spatial error leads to smaller corrections (5,9,10). This is either because one would weigh the final sensed error less compared to some internal prediction of the error, as predicted by Bayesian frameworks (11,12) (see Fig1A), or because the noise added downstream of movement planning (i.e. in movement execution and in the sensory signals about the error) is relatively large compared to the noise in planning (8). The possibility of online control while unfolding the action could also affect the relevance of the final temporal error. For instance, in open loop actions or when the movement time is very stable (e.g. the baseball example or in saccadic eye movements) the time of action onset determines the final temporal error (i.e. they are highly correlated) and one could weigh the final error less and base the corrections on some prediction error between the intended and actual action onset (Fig1A). This should be especially true for fast movements in which predictive components are crucial. Alternatively, both prediction and final errors might be used in combination to correct the next movement. We consider both possibilities in this study.

We know that predictions based on forward models (13) are important for correction mechanisms in general. That is, discrepancies between the prediction and some feedback, be it internal or sensed (14), are the key for mainstream computational models of motor learning (15,2) to explain the corrections of saccadic movements (16) or fast arm movements which are too brief to benefit from the final sensory feedback. In particular, in conditions where humans are aware of perturbations, learning from errors based on internal predictions can even override learning from final spatial errors (17) leading to the distinction between different kinds of errors: aiming errors (i.e. discrepancy between the planned and feedback positions) and target errors (i.e. target vs feedback position discrepancy), which are important in motor learning models (18).

Here, we resort to a similar distinction: errors based on the discrepancy between internally predicted and sensed action onset (prediction error) and temporal errors based on the experienced sensory feedback at the end of the movement. Because prediction error is more relevant in faster movements than in slower movements as the reliability of the prediction decays with time (13), we expect a different contribution of each error type in the next trial correction depending on how fast the movements are. We test this hypothesis by probing temporal corrections in different interception tasks.

We will consider the situation in which errors arise when inappropriate motor commands are issued (execution errors) as opposed to errors caused by external changes (19,5). In order to see the extent of the corrections, we exploit the properties of the time series of action onset in arm movements, saccadic eye movements and button-presses to study how people correct when the initial prediction error at action onset (see Fig1A) contributes differently to the final sensed temporal error with respect to a moving target in the different conditions. In the button press experiment, a keypress released a cursor to move with a fixed velocity profile to intercept the target. In this condition, as in the eye movements condition, the prediction error is highly correlated with the final temporal error (but see (16,20) for online saccade corrections). However, the final error can be perceived with high perceptual uncertainty in the eye movements condition due to the variability of saccadic reaction time and the previously reported temporal and spatial distortions at the time of saccades starting about 50 ms before saccade onset and continuing up to 50ms after saccade offset, a phenomenon often termed saccadic suppression (21,22). Finally, arm movements with different movement times will enable us to determine whether the relative contribution of each type of error depends on the movement time. A model based on a Kalman filter will be used to obtain an estimate of the predicted action onset and therefore, the prediction error. We show that prediction error relative to action onset and final temporal error relative to the target can be used in combination for trial-to-trial corrections. The contribution of each error signal has a specific time course following the action onset.

## Methods

### Arm movement experiment

#### Participants

15 subjects (age range 22-33 years, 11 males) participated in the experiment. Twelve of them were right-handed and three were left-handed as by self-report. All of them had normal or corrected-to-normal vision, and none had evident motor abnormalities. All subjects gave written informed consent. The study was approved by the local research ethics committee of the University of Barcelona (Institutional Review Board IRB 00003099). The experiment was conducted in accordance with the Code of Ethics of the World Medical Association (Declaration of Helsinki).

#### Apparatus

Participants sat in front of a graphics tablet (Calcomp DrawingTablet III 24240, 60 × 34 cm) that recorded movements of a hand-held stylus. Stimuli were projected from above by a Mitsubishi SD220U ceiling projector onto a horizontal back-projection screen positioned 40 cm above the tablet. Images were projected at a frame rate of 72 Hz and a resolution of 1024 by 768 pixels. A half-silvered mirror midway between the back-projection screen and the tablet reflected the images shown on the visual display giving participants the illusion that the display was in the same plane as the tablet. Lights between the mirror and the tablet allowed subjects to see the stylus in their hand. Virtual moving targets were white dots on a black background (shown red on white in Fig 1A). A custom program written in C and based on OpenGL controlled the presentation of the stimuli and registered the position of the stylus at 125 Hz. The software ran on a Macintosh Pro 2.6 GHz Quad-Core computer. The set-up was calibrated by aligning the position of the stylus with dots appearing on the screen, enabling us to present visual stimuli at any desired position of the tablet.

**Fig 1.**
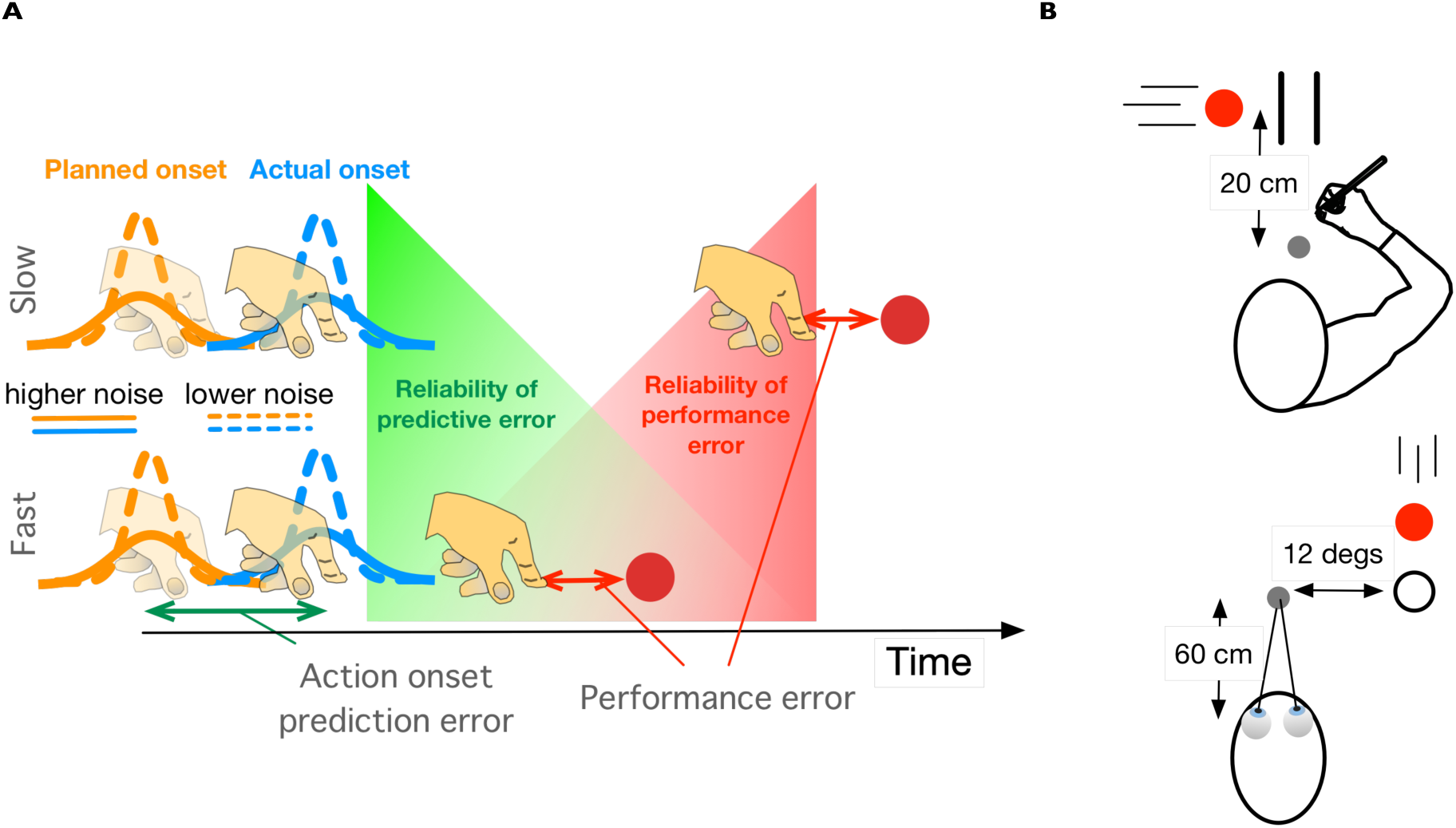
(A) Action onset and its reliability to predict the relevant task variable: temporal error with respect to the moving target. The uncertainty in determining the planning of the action onset (hidden variable) is illustrated by the orange Gaussians, while the execution noise is denoted by the blue Gaussians centered at the actual action onset. Different variabilities in the planning of action onset or its execution are denoted by the type of line (dashed: lower noise; solid: higher noise). The prediction error is the difference between the planned (or predicted) and actual action onset. The top row illustrates a slow movement after action onset (longer duration until crossing the target) and the bottom row a fast movement. One would expect larger corrections when the execution noise of the actual action onset is lower (blue dashed curves) relative to the planned noise (solid orange curves). The relevance of the prediction error is expected to decay after action onset (green area), while the increasing relevance of the final temporal error for planning next trial is denoted by the red area. (B) Illustration of two of the experimental tasks: arm movements (top) and eye movements (bottom).

#### Procedure

To start a trial, subjects had to move the stylus to the home position (grey dot in Fig 1B). After a random period between 0.8 and 1.2 seconds, a moving target that consisted of a white dot of 1.2 cm diameter appeared moving rightwards (or leftwards for left-handed subjects). Targets could move at one of three possible constant speeds (20, 25 or 30 cm/s), and were interleaved within the session. The target moved towards two vertical lines of 2 cm height and separated by 1.2 cm. The space between the lines was aligned with the home position (Fig 1B). Subjects had to hit the target (i.e. passing through it) at the moment the target was between the two vertical lines. Because we instructed participants to hit the target in the interception zone, we only had temporal errors associated to responses, except for the trials in which subjects missed the zone (less than 2%). The starting position of the target was determined such that it always took 0.8 s for the target to reach the interception zone. Auditory feedback was provided (100ms beep at 1000Hz) whenever the absolute temporal error between the hand and the target was smaller than 20 ms when the hand crossed the target’s path between the two lines. Each subject completed 360 trials (90 trials in 4 sessions).

#### Data analysis

The individual position data time series were digitally low-pass filtered with a bidirectional Butterworth filter (order 4, cut-off frequency of 8 Hz) for further analysis. Hand speed was computed from the filtered positional data by three-point central difference calculation. For each trial, we then computed the time of arm movement onset, the peak velocity, the movement time (elapsed time from the hand movement onset until the hand crossed the target’s path), and the temporal error with respect to the target. Movement onset was computed offline by using the A algorithm reported in (23) on the hand speed. This algorithm (a) locates the temporal location at which the velocity is greater than or equal to a tolerance value which is a percentage (10% in our case) of the peak velocity, (b) proceeds iteratively from this location backwards updating the tolerance value (10% of the current velocity sample) until the value of the velocity is less than the tolerance value.

### Button press experiment

#### Participants

Eight participants (age range 23-32 years, 5 males) participated in this experiment. All of them had normal or corrected-to-normal vision, and none had evident motor abnormalities. All subjects were right handed and gave written informed consent. The study was approved by the local research ethics committee of the University of Barcelona (Institutional Review Board IRB 00003099). The experiment was conducted in accordance with the Code of Ethics of the World Medical Association (Declaration of Helsinki)

#### Apparatus

Stimuli were shown on a Philips CRT-22 inch (Brilliance 202P4) monitor at a frame rate of 120 Hz and a resolution of 1024 by 768 pixels. The viewing distance was about 60cm and the head was free to move. A custom program written in C and based on OpenGL controlled the presentation of the stimuli and registered the time of the button-presses by sampling an ancillary device at 125 Hz. The software was run on a Macintosh Pro 2.6 GHz Quad-Core computer.

#### Procedure

The stimuli were the same as in the Arm Movement experiment except for the fact that the motion was presented on the fronto-parallel plane. In this experiment subjects had to press a button that initiated the release of a moving cursor from the home position. Subjects had to press the button timely so that the cursor would hit the target when passing between the two vertical lines (interception zone). The movement time of the cursor from the home position to the interception zone was always 330 ms and its velocity profile was extracted from an actual arm movement and was the same in every trial. This particular movement time was chosen because it is very close to the mean movement time of the fastest participant in the Arm movement experiment (325 ms) and would therefore limit possible online corrections. In this experiment the time of the button press determined completely the final temporal error. Subjects took the same number of trials and sessions as in the Arm movement experiment.

### Eye Movement experiments

#### Participants

Fifteen participants (age range 18–47, 7 males, including two authors) participated in the experiments. Among them, ten (age range 18–46, 4 males) participated in the first experiment (termed knowledge of results, KR) and twelve (age range 23-47, 5 males) participated in the second one (knowledge of performance, KP). They had normal or corrected-to-normal vision. All participants gave written informed consent. The study was approved by the local research ethics committee of the University of Barcelona (Institutional Review Board IRB 00003099). The experiment was conducted in accordance with the Code of Ethics of the World Medical Association (Declaration of Helsinki).

#### Apparatus

Stimuli were generated using the Psychophysics Toolbox extensions for Matlab^®^ (24,25) and displayed on a video monitor (Iiyama HM204DT, 100 Hz, 22’’). Participants were seated on an adjustable stool in a darkened, quiet room, facing the center of the computer screen at a viewing distance of 60 cm. To minimize measurement errors, the participant’s head movements were restrained using a chin and forehead rest, so that the eyes in primary gaze position were directed toward the center of the screen. Viewing was binocular, but only the right eye position was recorded in both the vertical and horizontal axes. Eye movements were measured continuously with an infra-red video-based eye tracking system (Eyelink II^®^, SR Research Ltd., 2000 Hz) and data were transferred, stored, and analyzed via programs written in Matlab^®^. The fixation point that was used as a home position for the gaze was a 0.4 deg×0.4 deg square always presented on the bottom left quadrant of the screen. The target was a 0.4 deg of diameter disk, and the interception area was a 0.6 deg of diameter ring. The interception area was located 12 deg to the right of the home position (see Fig 1B). All stimuli were light grey (16 cd/m^2^ luminance) displayed against a dark grey background (1.78 cd/m^2^ luminance). Before each experimental session, the eye tracker was calibrated by having the participant fixate a set of thirteen fixed locations distributed across the screen. During the experiment the subject had to look at the center of the screen for a one-point drift check every fifty trials. If there was any gaze drift, the eye tracker was calibrated again.

#### Procedure

A session consisted of 390 trials lasting between 2 and 2.45 secs. Each trial started with the subject looking at the fixation point for a period randomly varying between 700 and 1100ms. Participants were instructed to make a saccade to intercept the target, that was moving downward towards the interception area, at the time it was within the interception area. Targets moved with a constant velocity of either 20, 25 or 30 deg/s. Participants performed two versions of the KP experiment: KP-interleaved and KP-blocked. Target velocities were interleaved across trials in both the KR and KP-interleaved experiments. In the KP-blocked condition, each target velocity was presented in a separate 130-trial block, where the three blocks followed each other in a pseudo-random order counterbalanced across participants. The same participants experienced both KP conditions; the order was counterbalanced across subjects. The time to contact the interception area was 600 ms after target onset, and the target starting point was therefore depended on the actual target velocity. This duration (600 ms) was shorter than in the hand movements condition in order to adequate the target motion trajectory to a vertical movement in the screen. The occurrence of a saccade was crudely detected when the online eye velocity successively exceeded a fixed threshold of 74 deg/s. If the offset of an ongoing saccade was detected before the target reached the interception zone the target was extinguished at the next frame, i.e. within the next 10ms (offline analysis revealed that the target disappeared on average 2 ms after the time of the actual saccade offset). If the target center was aligned with the interception area before a saccade was detected we extinguished the target. Therefore, participants never saw the target after it had reached the interception zone. We delivered an auditory feedback (100 ms beep at 1000 Hz) if the eye landed within 3 deg of the interception area with an absolute temporal error smaller than 20 ms. To this end, the actual saccade onset- and offset-time and position were computed immediately after the saccade using the real-time Eyelink algorithm with a 30 *deg*/*s* velocity and 8000 *deg*/*s*^2^ acceleration thresholds (on average we retrieved these values 12 ms after the end of the saccade). In the first experiment (KR), participants did not receive explicit feedback on their performance other than the auditory one. In the second experiment (KP), the actual temporal error was displayed numerically in milliseconds at the end of each trial (KP). For offline analyses, a human observer validated each saccade manually. Saccades with an amplitude gain smaller than 0.5 or a duration longer than 100 ms were discarded.

### Analysis

#### Testing for the optimality of corrections: autocorrelation analysis

It is known that the serial dependence of consecutive movement errors depends on the amount of trial-by-trial correction (26). Suppose that no corrections of the performance error are made whatsoever (learning rate or correction fraction *β* = 0). In this case, we expect that final temporal errors will be similar to the previous one. The absence of correction would be revealed by a positive lag-1 autocorrelation function (acf(1)) of the performance errors under the assumption that planning noise accumulates from trial to trial as found in spatial corrections (8). The same prediction would be expected for the action onset, if it determines completely the final temporal error (e.g. Button press experiment). On the contrary, if one aims at correcting for the full observed error (*β*=1) then consecutive movements will tend to be on opposite sides of the average response because one corrects not only for the error in planning but also for the random effects of execution noise and sensory noise. In both scenarios (*β*=0 and *β*=1) there is an unnecessarily large temporal variability, but this has different causes. If one does not correct, not only will previously committed errors persist but also previous planning errors will accumulate across trials increasing the variability much like when one repeatedly reaches out for static targets. If one does fully correct, the variability due to changes in the planned time will be larger than if smaller corrections were made. In either case the process is not optimal in the sense that the temporal error is more variable than necessary. When corrections are large enough to compensate for planning errors but not too large to systematically overcompensate for planning errors, then the temporal error variance is minimal and the correction fraction is optimal. For such fractions of corrections, acf(1) of the temporal errors will be zero (8). We conducted simulations (see below) showing that optimal corrections that minimize the final temporal variability also produce zero acf(1).

#### Dependency on the previous final temporal error

We analyzed the dependency of the time of action onset in the current trial on the finaltemporal error with respect to the target in the previous trial in the different conditions by fitting linear mixed-effect models (LMMs), which enable us to analyze the effects of the previous trial on the current response. Since we expect that the effect of the previous final temporal error on the current action onset could be modulated by the presence of online corrections and the type of movement (fast or slow) we considered these variables in the model. As an indicator of online corrections we computed the number of peaks in the velocity profile of the arm movements and the number of corrective saccades (i.e. saccades occurring within 400 ms after the primary saccade offset and lending in the vicinity of the interception area) in the saccade experiments. Note that this cannot be applied to the button press. Within each participant we also categorized the movement in every trial as slow or fast depending on whether the duration was shorter (fast) or longer (slow) than the median movement time of the participant.

In the model, the action onset time was the dependent variable and the final temporal error and the number of velocity peaks in the previous trial were the independent variables (fixed effects). Participant and type of movement in the previous trial (fast or slow) were included as random grouping factors. Both intercept and slope were allowed to vary as random effects. We used the lmer function (v.1.0–6) (27) from R software.

As illustrated in Fig1A, we expect a larger contribution of the final (target) temporal error for slower movements in the Arm movements experiment and less contribution in the open loop conditions (Button press and Eye movement experiment). However, for slower arm movements we expect this contribution to be reduced when online corrections were present in the previous movement.

#### Simulations and modelling of temporal corrections

We will implement the temporal corrections at the action onset across simulated trials in which we will manipulate different sources of variability: variability in planning and execution variability. In each simulated trial *i* the generation of a planned action onset *τ* is a stochastic process where *τ*_*i*_, the planned action onset at trial *i*, is updated according to:

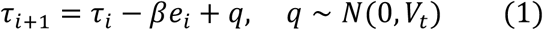

where *β* is a learning rate or, in our case, the simulated fraction of correction with respect to the final temporal error (*e*) and *q* is the process or planning noise which we assume is Gaussian noise with variance *V*_*t*_. The actual action onset *T*^*s*^ is simulated by adding execution noise (produced by execution noise) to the planned action onset:

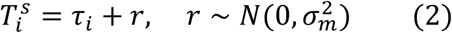

where *r* is the execution noise (added noise from when the motor command is issued until movement onset). The final temporal error *e* at trial *i* is given by:

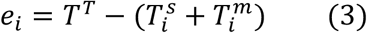

where *T*^*T*^ is the time at which the target is centred within the interception zone and 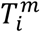 is the movement time. Without loss of generality, we set 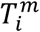 and *T*^*T*^ to zero in the simulations.

The planning of the time of action onset to intercept a moving object can be affected by different sources of noise (*V*_*t*_ in eq. 1) or temporal variability (28) that can be expressed as follows:

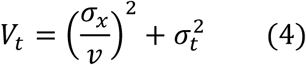

The first term is velocity dependent and the second one corresponds to a timing variability (28), (29). *σ*_*X*_ is the spatial variability about the target position at action onset and *ν* is the target speed. Uncertainty caused by measuring target speed may likely contribute to the timing or velocity-dependent variability. However, in practice both sources of variability are difficult to tease apart because an error in misjudging the target position would be indistinguishable from a timing error (28). We therefore do not distinguish between these different sources but treat them as a single noise source.

##### Modeling the corrections

Using the equations introduced above, we modeled a trial-to-trial correction of the time of initiation, assuming that all the final temporal error is fully caused by the time of action initiation *T*^*s*^. This was certainly the case in the Button press experiment – because in our case the time to reach the target was fixed once the button was triggered - and to a lesser extent in Eye movement experiment, while for arm movements there is some room for online corrections by adjusting the movement time. We modeled 16 different correction fractions from 0.06 to 0.96 with increments of 0.06 (range: 0.06-0.96) and four values of *r* (*σ*_*m*_ = 0.022, 0.05 0.1 and 0.2 s). We set *σ*_*x*_ to 1 cm and *σ*_*t*_ to 0.05 s. These values were used with three target velocities: 20, 25 and 30 cm/s resulting in a mean process noise variance of 0.0042 s2. These choices were guided by values reported in previous studies (28,30). If the simulated time at trial *i* was shorter (i.e responding too early) than a target value (e.g. 0 ms) by some magnitude *e*_*i*_, the value of the intended time onset (*τ*_*i*+1_) was increased by *βe* on the next trial, or decreased if the observed time was too long. We ran 1000 simulations for each combination of *β* and *r*. Each simulation consisted of a series of 360 responses or trials in which speed was interleaved (but note that the time the target took to reach the interception zone was the same for all speeds, so target speed changes between consecutive trials do not prevent trial-by-trial corrections).

In the simulations we know the (simulated) internal state (i.e. planned action onset *τ*) and both process and measurement noise. A Kalman filter model would basically estimate this internal state from the two types of noise. In addition to estimate the planned action onset, the Kalman filter will also provide us with a sort of learning rate (Kalman gain *K*) that optimally corrects the planned time as a function of the temporal variance of the planning process *V*_*t*_ and the variance of the observation or measurement 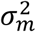 (5):

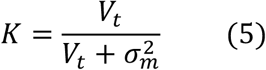

This way we will know whether the corresponding Kalman gain in our different simulations matches the fraction of correction (*β*) that minimizes the temporal variance (i.e. optimal correction) and produces an acf(1) near zero.

#### The Kalman filter model on the behavioral data

Unlike in the simulations, we do not know the planned action onset in the behavioral data. In order to estimate the prediction error of the action onset we need to estimate the planned action onset (*τ*_*i*_) in each trial *i*. We then applied the Kalman filter model to estimate the planned action onset and determine the degree of correction based on the prediction error. As shown in eq. 2 the actual action onset *T*^*s*^ is a noisy realization of the predicted action onset *τ*. We can rewrite eq. 1 as:

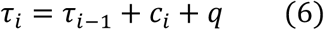

where *c*_*i*_ is a correction factor that has to be determined by the Kalman filter. But, how does the Kalman filter work out the magnitude of the correction? The Kalman filter estimates *c*_*i*_ recursively by combining a predicted or planned action onset (i.e. a priori) and the observation of action onset *T*^*s*^ that has been corrupted by noise. After movement onset at trial *i*, the Kalman filter estimates a posterior time of action onset (denoted by the hat operator):

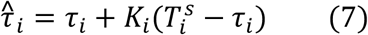

The posterior will be used as a predicted action onset time in trial *i+1*, becoming *τ*_*i*_ in (eq. 6). *K*_*i*_ is called the Kalman gain and reflects the fraction of correction of the prior time of action onset. If *K* = 0 no change is made in the planning for the next trial; alternatively, if *K* = 1 the whole difference between the prediction and the observed action onset is accounted for in the posterior. We will refer to the difference between *T*^*s*^ and *τ* as *prediction error*.

In order to compute *K*, the Kalman filter takes into account the uncertainty of the prediction and of the observation.

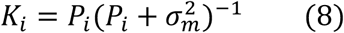

where *P*_*i*_ is the uncertainty (the variance) in the prediction of the planned onset time before the observation of action onset takes place. Note the equivalence with eq. 5. This *a priori* uncertainty is obtained from the posterior estimate of the uncertainty,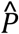, in trial *i-1*:

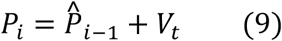

The Kalman filter will correct the internal estimate (i.e. predicted action onset) by a fraction *K* of the prediction error *T*^*s*^ − *τ*. However, although the prediction error is highly correlated with the final temporal target error in some conditions, the prediction error is not the task-relevant error shown in eq. 3. We analysed the correction with respect to action onset because we are interested in how people correct in the planning phase.

The planning of the action onset should aim at minimizing the expected final temporal error (*e*(*τ*) = 0) which can be stated as:

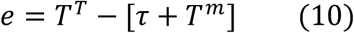

In order to be accurate across all observed responses we need that:

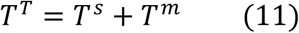

Substituting eq. 11 in eq. 10:

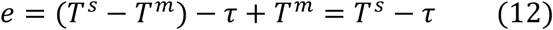

which is the prediction error with respect to action onset that the Kalman filter corrects for. This equation shows that, given some constraints on the distribution of movement time *T*^*m*^ (i.e. the same distribution as *T*^*s*^ with the mean shifted), correcting for the prediction error is equivalent to correcting for the final temporal error. This is true on average only, since for individual trials the prediction error does not necessarily correspond to the final error and because the prediction error is independent of *r*, but the final temporal error is. Note that we have used *β* as the learning rate with respect to the final temporal error and *K* (Kalman gain) as the learning rate with respect to the predictive error.

##### Parameter estimation

In order to estimate the predicted action onset time (*τ*)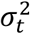, the variance of the execution noise was the only free parameter as it is usually the case in Kalman filter models (31). 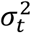 was determined by least-squares optimization and the squared error values were computed on d Kalman filter modelsifferences between model prediction of the planned onset time and actual observations of the action onset on the same trials. The process variance (*V*_*t*_) was computed from the variance of the observed time of action onset by using a moving window of 4 trials as in (31).

### Data availability statement

Data that support the findings of this study have been deposited in the open science framework and can be obtained in the following link: https://osf.io/u273g/?view_only=c32ecbc3b4c24f9d8f1354d0b2d6ee50

## Results

We show in Fig 2A examples of series of observed action onset times from the different experiments. From the different series we then computed the acf(1) for both the action onset and final temporal error as temporal markers. To qualify as “optimal correction”, acf(1) of target error must be not-significantly different from zero. Fig 2B shows the mean acf(1) across subjects for the two temporal markers in the different experiments. As can be seen the acf(1) for the two moments of interest are virtually the same in the Button press and eye movements experiments. This is expected and basically confirms the high dependency between action onset and final temporal error and the fact that the time of action onset can be used to correct the performance error. Except for the action onset in the arm movements experiment, the 95% confidence intervals of the acf(1) include the non-significant interval (horizontal grey lines). The only acf(1) value whose 95% confidence interval is well above the significant level is the action onset of the arm movements. Furthermore the difference in acf(1) between the action onset and final temporal error, regarding the acf(1) can be interpreted as the action onset of arm movements not being entirely responsible for correcting for the final temporal error. This could mean that online corrections during a movement might have contributed to achieve a closer to zero acf(1) for arm movements. It is known that people can adjust the timing of a movement online (without changing the spatial trajectory) with latencies of 169 ms (32). Therefore these adjustments can account for people achieving near optimal corrections of the final temporal error while making under-corrections of the action onset (positive significant acf(1)). For the other conditions, we can see that final temporal errors were corrected in a near optimal way.

**Fig 2.**
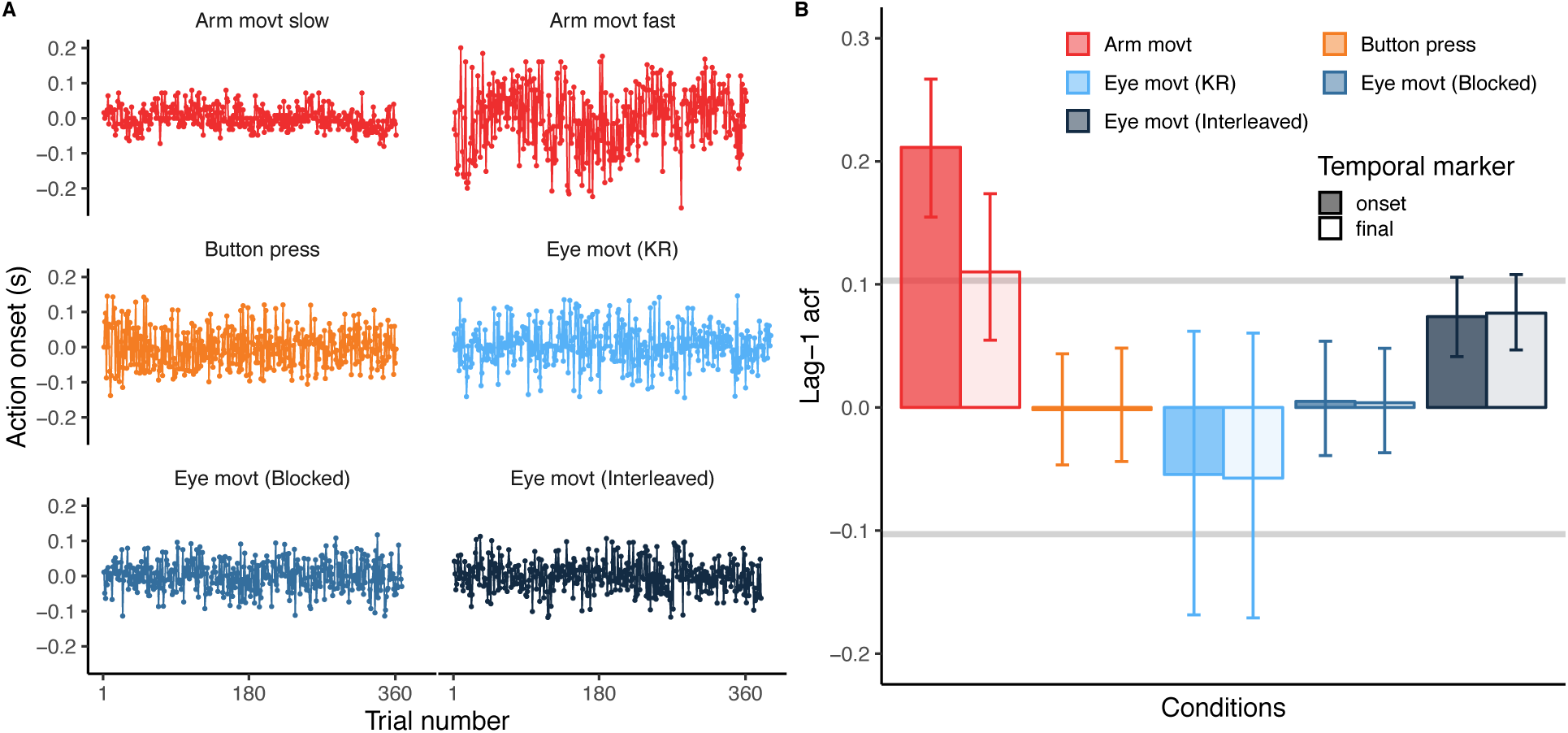
(A) Example of action onset times for the different experiments in each panel. Different experiments are color-coded (the same as panel B). Each response series corresponds to a single participant. The two examples of the Arm movement experiment correspond to a slow (top-left) and fast (top-right) participant. The action onset time is centered at zero (by substracting the mean) to optimize panel space. (B) Mean lag-1 auto-correlation functions, acf(1) for the time of action onset (solid bars) and final temporal error (empty bars) for the different conditions. Error bars denote the 95%-CI of the correlation coefficients. The two grey horizontal bars denote the significance of the acf.

In order to provide further evidence that a near zero acf(1) minimizes the temporal variance, we looked next at the results obtained from the simulated data. Fig 3A shows how the simulated fraction of correction with respect to the final temporal error modulates the overall temporal variance of this error. The correction fraction with respect to the final temporal error (*β*) for which the temporal variance is minimal is the optimal correction fraction. As can be seen, this fraction is different for the different levels of simulated measurement noise (*r* in eq. 2) that correspond to the different Kalman gains: the larger the simulated measurement noise the smaller the optimal correction fraction or gain eq. 5. Importantly when we plot the acf(1) of the temporal errors against the fraction of correction (Fig 3B) we can see that the values corresponding to optimal corrections in Fig 3A produce acf(1) values that are very close to zero. From the different data patterns shown in Fig 3 we can be quite confident that the Kalman gain denotes a correction fraction that is optimal in our simulated temporal correction task.

**Fig 3.**
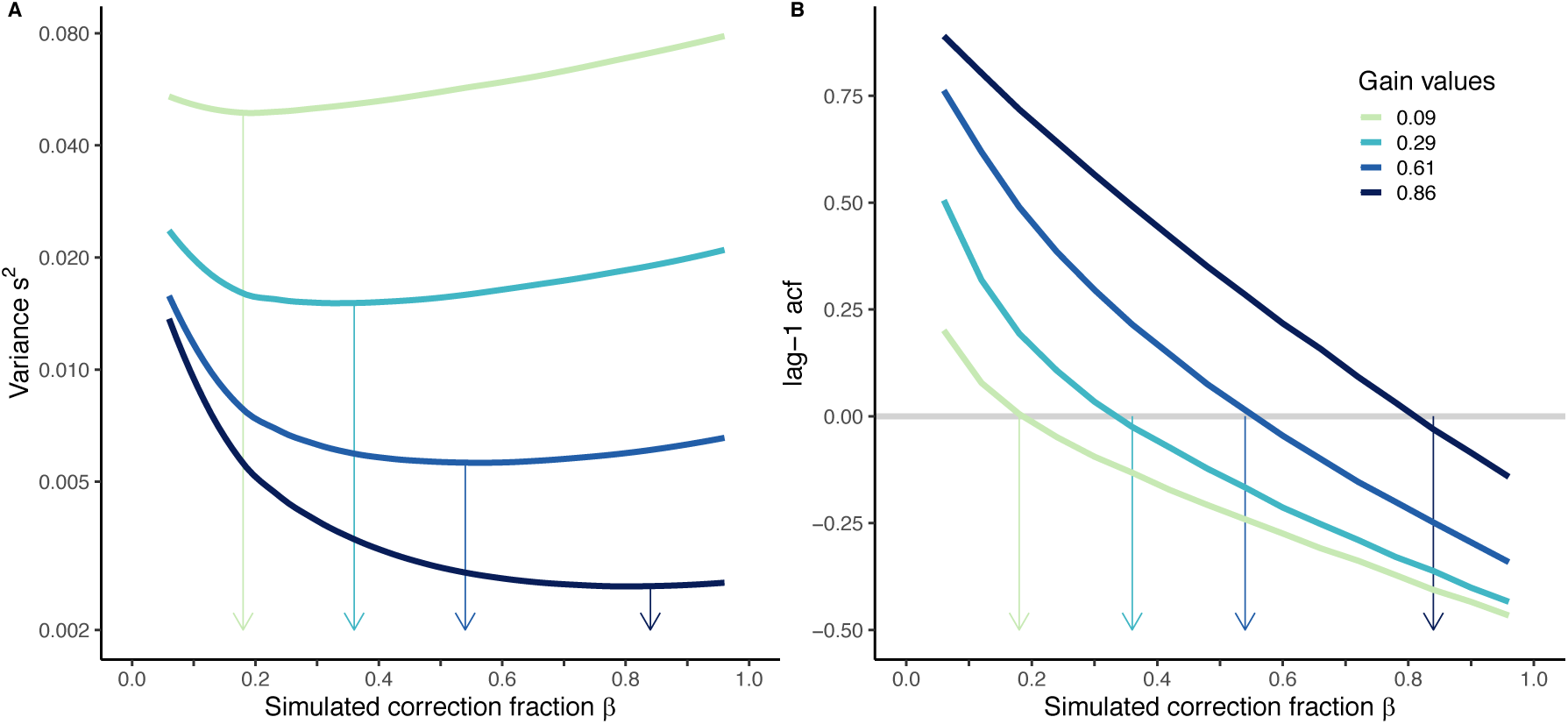
Simulation results. (A) The temporal error variance as a function of the simulated fraction of correction β for the four different levels of simulated execution noise (from bottom to top:σ_m_= 0.022, 0.05 0.1 and 0.2 s). The arrows point to the β value corresponding to the minimum variance. As can be noted, this fraction of correction is similar to the simulated gain (see legend in panel B, the gain in turn depends on the level of execution noise). The largest gain (i.e. K=0.86) requires larger corrections in order to minimize the variance. (B) The acf(1) values of action onset in the simulated data against the correction fraction. As can be seen, the acf(1) should be near zero for each gain to be optimal for minimizing the variance.

### Are people making temporal corrections?

Fig2B shows a specific pattern of correction. However, one concern is whether people were actually making temporal corrections. The task design encourages the use of the final temporal error as the task-relevant performance error (32). This is achieved by designating a fixed interception zone, so that participants can, in terms of a systematic error, only be either early or late with respect to the moment at which the target is at the center of the interception area. Although it is always difficult to separate temporal from spatial errors, in addition to the linear models described above which contains the previous temporal error as fixed effect, we also ran the same model with the previous spatial error as fixed effect and used the Akaike Information Criterion (AIC) to compare the corresponding models. The AIC for the model in which the previous temporal error was specified as fixed effect was smaller than the AIC for the spatial error model (−47370.03 vs −47338.19) denoting a better description of the previous temporal error of the action onset in the following trial.

Assuming that open-loop control schemes are used to execute the movements, we expect (i) a modulation of the initiation times by prior temporal errors and (ii) that the time of action initiation relative to the interception time does not statistically differ depending on target velocities. That is, relevant decision variables regarding the action onset would mainly rely on temporal estimates of the remaining time to contact from the action initiation. An ANOVA on the linear mixed model in which action onset was the dependent variable, target speed (fixed effect as continuous variable), conditions (fixed effect as factor) and subjects treated as random effects failed to report a significant effect of target speed on action onset (F<1, p=0.96) and only condition was significant (F=53, p<0.001). The interaction was not significant (F<1, p=0.473). Overall, these results reinforce the assumptions that subjects were using temporal errors rather than spatial errors and that the action onset was mainly driven by the time to contact information.

### Dependency on the previous temporal error

At this point we cannot tell how the observed corrections were made: participants could have changed their action onset with respect to some prediction error or with respect to the final temporal error.

Autocorrelation indicates how consecutive points tend to be around the mean (e.g. if one overcorrects then consecutive points will likely be on opposite sides), but does not indicate which fraction of the previous final error is being accounted for in the change of action onset in the present response. In order to get an estimate of this magnitude we ran the Linear Mixed Model (described in the Methods sections). The time of action initiation at each trial was fitted as a function of the previous final temporal error. The slope denotes how much the previous error is considered. Fig 4A (red dots for the Arm movement experiment and boxplot) shows the values of the slopes. The only significant slope (slope=0.18 fraction/trial, t=5.56, p<0.0001) was found in the Arm movement experiment (see inset in Fig 4A). The slope was 0.01 (t=-0.432, p=0.688), 0.05 (t=0.916, p=0.36), −0.037 (t=0.829, p=0.41), 0.05 (t=1.330, p=0.184) for the Button press, Eye movement (KR), Eye movements (Blocked) and Eye movements (Interleaved) experiment respectively.

**Fig 4.**
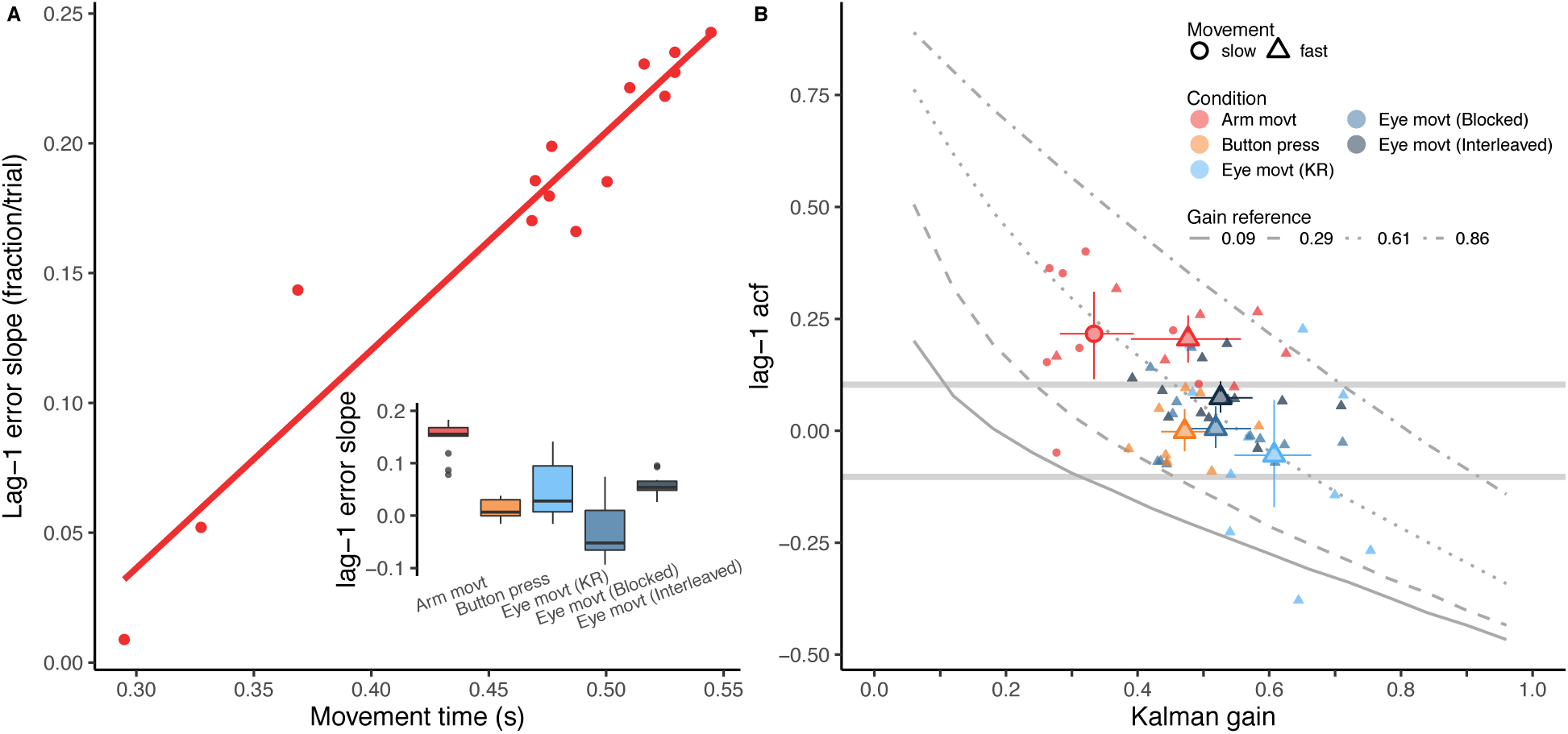
(A) Dependency (slope in the linear model) of the action onset on the previous final temporal error as a function of movement time in the Arm movement experiment. Each dot corresponds to an individual participant. (inset) The boxplots show the mean slopes of all the experiments. (B) The acf(1) for each participant against the estimated Kalman gain (K). Smaller symbols correspond to individual participants and conditions, while larger symbols are mean values across subjects within conditions. For the Arm movement experiment we split the data points into slow and fast participants depending on the movement time (shape coded). Error bars denote 95%-CI. The two horizontal grey curves denote the confidence interval for the null acf(1). For the sake of comparison, the four lines with different styles (solid, dashed, dotted and dash-dotted) correspond to the Kalman gain and expected acf obtained in the simulations (see Fig 2B).

Since, the slope is only significant in the Arm movement experiment, we plot the individual slopes in the main panel of Fig 4A as a function of the mean movement duration of the arm movement. Interestingly, this reveals a clear positive linear relation between the movement time and the dependency on the previous temporal error. Participants with slower arm movements modified the action onset in the present trial more as a function of the previous final temporal error than participants with faster movements. However, this effect could be modulated by online corrections defined by the number of velocity peaks corresponding to secondary corrective movements, which was included in the model. The percentage of trials in which more than one peak (or one saccade) was observed was 38.8, 28.8, 27 and 27 respectively for the Arm movement, Button press, Eye movement (KR), Eye movements (Blocked) and Eye movements (Interleaved) experiments. This factor reduced the dependency of action onset on the previous temporal error as expected in the Arm movements experiment: each peak reduced the slope, and then the contribution of the temporal error, by 0.02, however this interaction was not significant (p=0.17) and neither was it for the Eye movement experiments (KR, Blocked and Interleaved): p=0.679, p=0.31, p=0.7. Note that this analysis was not possible in the Button press experiment.

The question then is how do people correct in these conditions where the effect of the previous temporal error is not significant (slope not different from zero)? One possibility (depicted in Fig 1A) is that people corrected the planned action onset, based on the difference between the planned action onset and the actual action onset (i.e. the prediction error) rather than on the final temporal error. Since we could not measure this prediction error in the experiment, we had to model correcting based on this error to infer how large these corrections were. As explained above, we used a Kalman filter model to estimate the Kalman gain, that is the fraction of the prediction error that is used to update the planned action onset for the next trial.

### Corrections based on the action onset prediction error

We fitted the Kalman filter as described in the Methods sections. The Kalman filter was fitted to the time series based on the action onset with the measurement noise variance 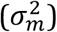 as the only free parameter.

The individual as well as the mean Kalman gains for participants and conditions are shown in Fig 4B together with the corresponding value of the acf(1) of the action onset. We chose to plot the acf(1) of the action onset because the Kalman gain reflects the correction of this temporal variable with respect to the prediction error. This plot shows that different values of correction with respect to the action onset prediction error (i.e. Kalman gain) can correspond to optimal or near optimal corrections for the Button press and Eye movement experiment. Interestingly the values of the Kalman gain are larger for those conditions in which the dependency on the previous final temporal error is non-significant: Button press and Eye movement and the faster arm movements, which show less dependency on the final temporal error (Fig 4A). This means that there is a larger fraction of correction (between 0.45 and 0.6) of the planned onset time with respect to the prediction error (action onset minus planned onset) in these conditions than for the slower arm movements (Kalman gain of 0.33).

A one-way ANOVA on the Kalman gain with condition as independent variable (separating slow and fast arm movements) yielded a significant effect of condition (F=8.36, p<0.0001). Multiple post-hoc comparisons revealed statistically significant differences between the Kalman gain in the slow arm movements and the rest of conditions (corrected p-values using False Discovery Rate (33) were p=0.016 with fast arm movements, p=0.002 for Button press, and p<0.0001 for the remaining conditions. The gain in the Eye movements (KR) was also significantly different than the gain in the fast arm movements experiment (p=0.04) and Button press (p=0.04).

The mean values for the different conditions, especially for the Button press and Eye movement experiments, cluster around one of the gain reference lines (k=0.61) obtained in the simulations. This line corresponds to a simulated measurement noise (r in eq. 2) of 50 ms, while the other simulated values were 22 ms (gain 0.86), 100 ms (gain 0.29) and 200 ms (gain 0.09). The estimated magnitude of the measurement noise SD were 36 ms (slow arm movements), 63 (fast arm movements), 76 ms (Button press), 75 ms (Eye movements KR), 77 ms (Eye movement blocked) and 76 ms (Eye movement interleaved). Except for the lower measurement noise in the slow arm movements, the estimates in the remaining conditions were not significantly different from 50 ms. We reasoned that, given the similar value of measurement noise across experiments, the lower value of the Kalman gain in the slow arm movements condition might be attributed to the lower process noise in this condition.

Higher Kalman gains can be interpreted as changing the a priori planned action onset by larger fractions of the prediction error (i.e. difference between the planned action onset and the actual movement onset). This would certainly be expected in the Eye movement experiments because the sensory feedback of the final temporal error can be noisy. However this is not the case for the Button press experiment in which participants could perfectly perceive the error. Since the final temporal error is fully explained by the time of action onset in this condition, it seems that the correction based on the prediction error rather than the final error is based on the reliability (or correlation) of the prediction error with respect to the final temporal error.

### Relation between prediction error and final temporal error

The previous analysis on the Kalman gain produced a significant difference between slow and fast arm movements: faster subjects corrected the planned action onset more (0.48 vs 0.33) on the basis of the prediction error. In addition, an opposite trend was found with respect to the dependency of the action onset on the final temporal error (see lag-1 slope shown in Fig 4A): slower subjects (longer movement times) corrected the action onset more with respect to this final error. To have a closer look at these two different trends we plotted both the obtained Kalman gain and the slope together as a function of the movement time in Fig 5. As can be seen, the relevance of the prediction error denoted by the Kalman gain decreases in a non-linear manner, while the contribution of the final target error (feedback-based weight) increases approximately linear. Moreover, Fig 5 shows that the contribution of the final temporal error starts to increase after a movement time or sensorimotor delay *δ*_*sm*_ (close to 280 ms). Is there a possible model that simultaneously describes the two trends such that by increasing one contribution (i.e. *β*) the other (*K*) decreases in a coupled way? It is clear that *β* and *K* are not linearly related, for example like in *K*_*t*_ = *c* − *β*_*t*_, where *c* would be the maximum fraction of correction found in the Kalman filter model. The decay of *K*_*t*_ rather suggests a quadratic decay of *K*_*t*_ as follows:

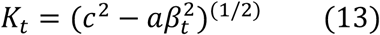

**Fig 5.**
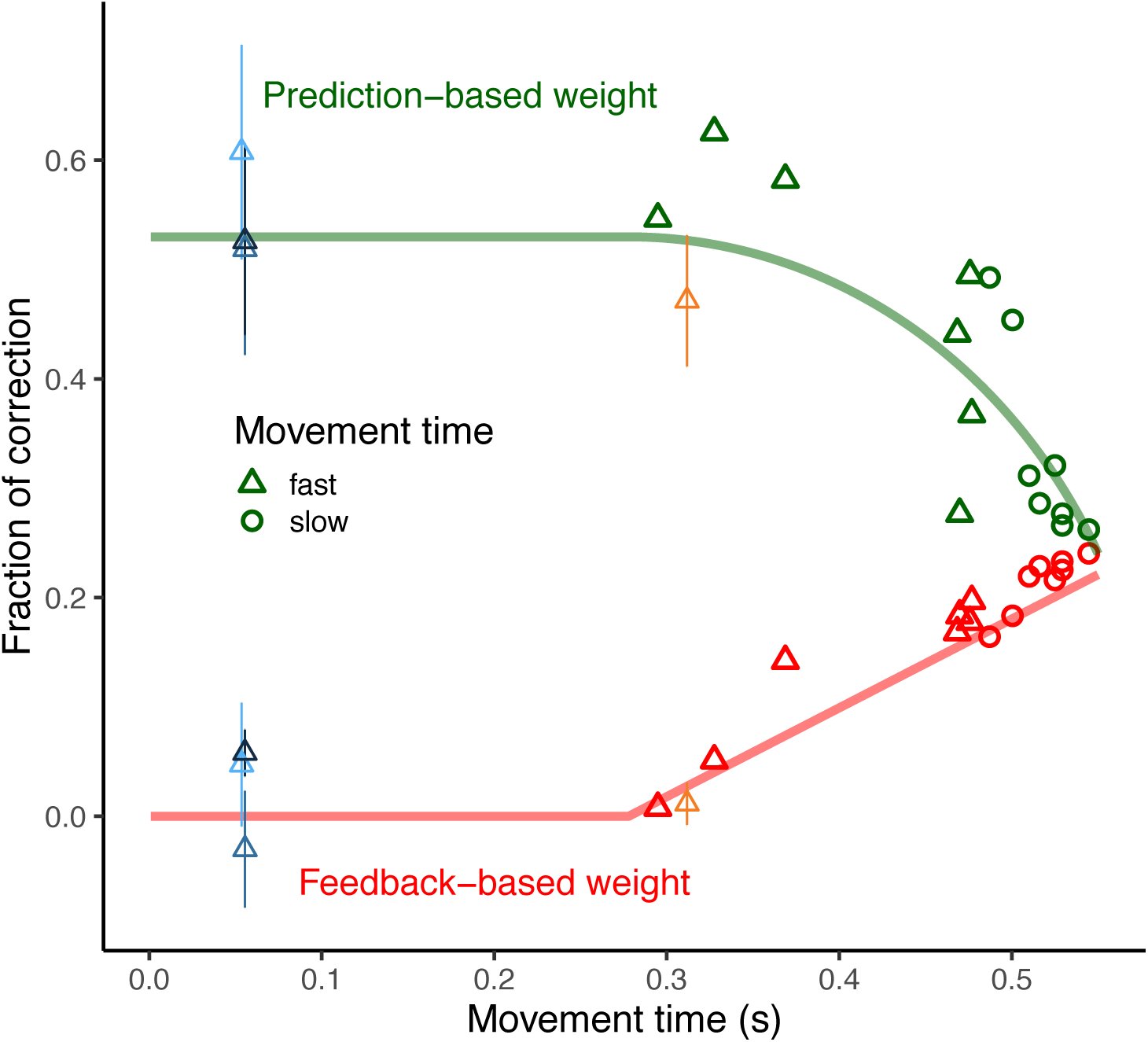
Evolution in time of the correction fractions (i.e. relevance given to prediction -green- and final error -red-) for the Arm movement experiment. Each point corresponds to a different participant and the symbol corresponds to the classification of the movement time duration (triangle : fast, shorter than 0.5 s; Circles: slow, longer than 0.5 s). The feedback-based weigh corresponds to the actual fraction of correction of the final error obtained from the slopes shown in Fig 4A using a linear model. This linear model was obtained using the simulated data and allowed to estimate the corresponding slope for each (simulated) fraction of corrections. The Kalman gain contribution is obtained by scaling the Kalman gain between 0 and 1. The shift of the two curves (prediction and feedback-based correction) depends only on the time at which the final temporal error will start to be considered for correction in the next trial (277 ms in the figure). See text for the computation of the two prediction curves. The mean values of the Kalman gain and the target error contribution (slopes) for the experiments other than the Arm movements are also plotted but not used in the fitting procedure. The color codes are the same as in Fig 4B.

We used eq. 13 to fit the decreasing Kalman gain and *β*_*t*_ (red line in Fig 5) was set in the same fit as a function of the movement time according to the following piecewise function:

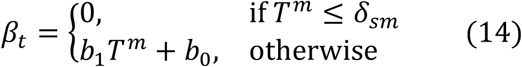

where *T*^*m*^ is the movement time and *δ*_*sm*_ the fitted delay after which the final performance error will be weighted for next trial correction. In order to couple the non-linear decrease with the linear increasing in the fitting procedure, we did not directly fit parameters *b*_0_ and *b*_1_ but rather restricted the value of *β*_*t*_ to (1 − *c*)^2^ at the movement time *T*^*m*^ = 550*ms*, so that the rate of increase of *β*_*t*_ depends on the parameter *c* also present in *K*_*t*_. Therefore, *δ*_*sm*_, *a* and *c* were the free parameters in the model plotted in Fig 5 so that the decrease of *K*_*t*_ and simultaneous increase of *β*_*t*_ minimized squared errors across the red and green symbols in Fig 5. The obtained values for these parameters were *δ*_*sm*_ = 277*ms, c* = 0.53 and *a* = 4.45

One alternative possibility is that people could use a combination of the two error signals (not contemplated in our model) resulting in some additional benefit than when using either signal alone. However, in our case, the prediction error and the final target error are highly correlated in most of our conditions (see Fig 6). Only in the case of the Arm movements experiment, the correlation between both error signals is low, about 0.25 (see bar plot in Fig 6). Due to the magnitude of this correlation, there is some small benefit in correcting based on integrating or combining both error signals (34).

**Fig 6.**
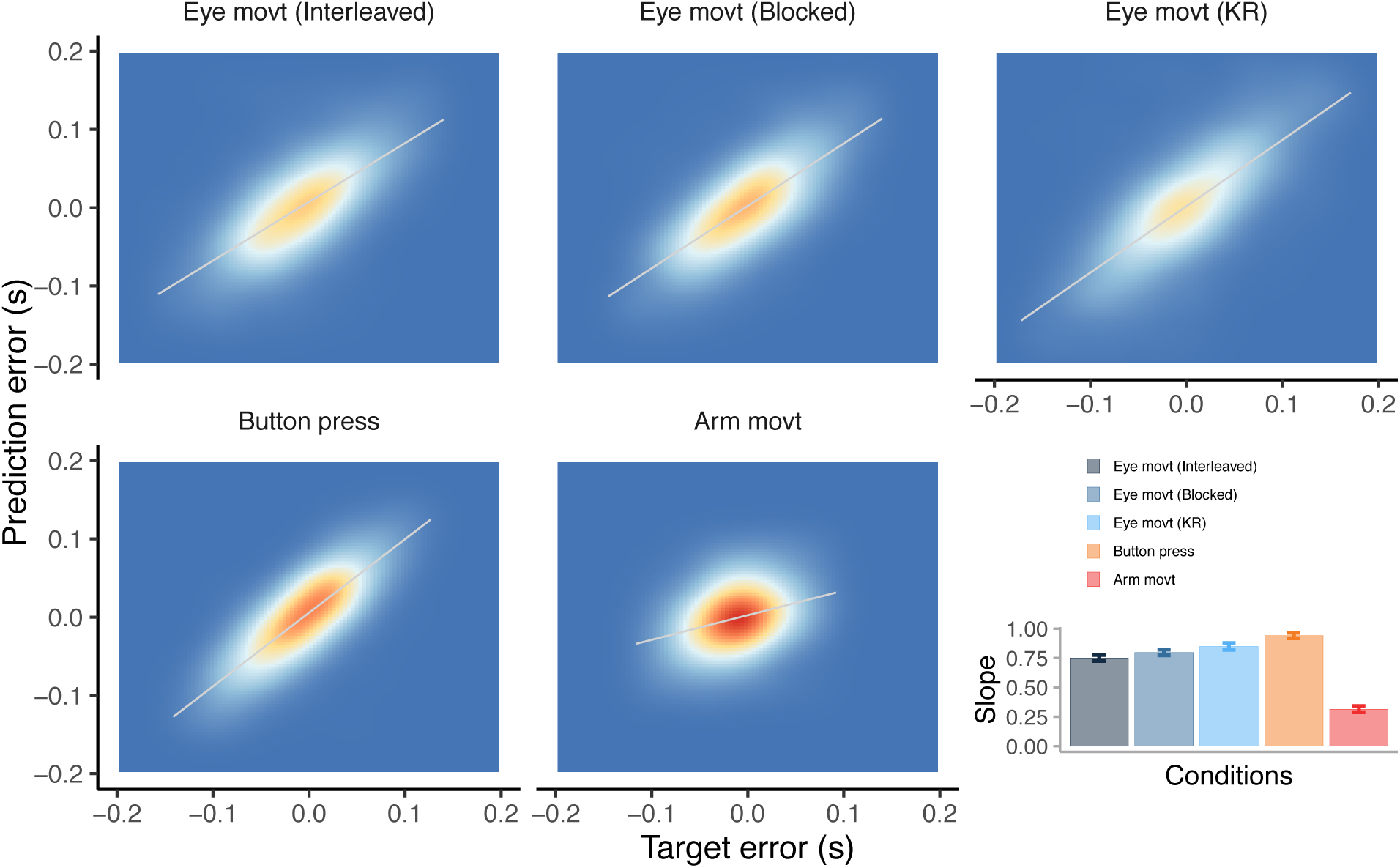
Joint density of the final temporal error (x-axis) and prediction error (y-axis) that is computed from the Kalman filter for each condition. The density plot includes all participants. The bar plot shows the slope of the fitted (grey) line for each condition. The slope for the Arm movement experiment is shown without separating fast and slow movements. However the slope was significantly smaller for slower movements (0.49 versus 0.72, p=0.01.)

## Discussion

We show that people minimize the temporal variance across trials when correcting for temporal errors. This is concluded from the structure of temporal correlations (35). We report non-significant, thus near zero, lag-1 acf of the previous temporal error in several interception tasks (see Fig 2B). However, which error signal is predominantly used to correct differs depending on the condition and duration of the movement of the action: slower movements showed larger dependency on the previous final temporal error with respect to the target in the Arm movement experiment. This dependency was modulated by possible online corrections. The more likely an online correction, the lesser the final error was considered for correcting next trial. In most conditions (Eye movement, Button press and fast arm movements), trial-to-trial correction based on the previous performance errors were not significant, i.e. lag-1 slope was not significantly different from zero (see inset Fig 4A). However, people rely on the prediction error at action onset rather than the final temporal error with respect to the target to change the planned initiation time in the next trial. This is based on the high values of the Kalman gain, which denotes that the prior (planned) action onset will be shifted in the next trial by a large fraction (between 0.45 and 0.6) of the prediction error (i.e. difference between the prior and actual action onset). In the Arm movements experiment, fast movements were initiated later, therefore their planning could have been more robust than slow movements (initiated earlier) increasing the reliability of the prediction error. The reliance on the prediction error has obvious advantages when the final sensory feedback is noisy. This is the case in the Eye movements experiments where perception of the sensory temporal error signal can be noisy. Correcting on the basis of the prediction error at action onset is possible under some restrictions (e.g. open loop), as there is a clear correlation between the prediction error and the final temporal error (Fig 6). Due to the possibility of making corrections during the movement, the correlation is lower in the Arm movement experiment, and this is the condition in which we find less contribution of the prediction error (slower movements).

Models of motor learning would predict less correction (e.g. lower learning rate) when the sensory feedback is more uncertain (5,10,11,9) or if error signals are perceived less relevant (7,36). The Bayesian explanation is that the sensory error feedback is weighted less in favor of internal state estimate (37). This is usually the case when studies focus on the reliability of the final task-relevant error. Our findings, however, show that the picture can be more complex. We found the same amount of correction in some of the Eye movement conditions and in the Button press experiment while the final temporal error signal is likely perceived with different uncertainty, as the Button press is more reliable. Our results show that corrections in these two conditions are executed in a very similar way (similar Kalman gains). The way temporal errors are corrected (mainly in the Button press condition) challenges some of the assumption of current models, and merely considering the final sensory error might not suffice, at least when temporal errors are relevant. The timing in the Button press experiment can only be controlled at action onset and this seems to be an important factor controlling which error signal will be used to correct in the next trial.

Prediction errors have been mostly regarded as relevant for online corrections when the sensory feedback would arrive too late to make useful corrections. Here we show that prediction errors can also be useful for trial-by-trial corrections. This is similar to previous findings that have shown the performance of predictive saccades in a given trial is affected by latencies in previous trials (38). The process of correcting a planned action onset apparently is not affected by low uncertainty of the final error (e.g. Button press experiment) because it does not override the use of the prediction error at action onset. Interestingly, it seems that the contribution of the final error for next trial correction depends on the movement time in arm movements. We found that the final target error will start to be weighted for correction in the next trial by movement times close to 277 ms. The model (Fig 5) also predicts that, as movement progresses, the reliability of the prediction error at action onset decreases reaching a minimum close to 600 ms. This time course of the contribution of the prediction error for the next trial parallels the shift from prediction to sensory signals in online correction of spatial errors of arm movements (13).

The evolution of the respective contribution of prediction and final errors suggests that the system has some access to or knowledge about the noise that is added from the time of action onset. This is in agreement with previous work showing that the motor system is able to model the temporal uncertainty of the movement time when programming reaching movements under temporal constraints (39).

The relevance of the prediction error in trial-to-trial temporal corrections is mostly noticeable in the Eye movements experiments. The behavioral plasticity of the saccadic system has been well established in the temporal domain: saccade latencies may be strongly affected by a number of factors such as temporal stimulus arrangement (40), stimulus properties (41,42), urgency (43), expectations (44) or reinforcement contingencies (45,46). Moreover, studying saccades directed toward a moving target revealed that the saccadic system takes into account both the saccade latency and duration, and is able to adjust to experimentally induced perturbations (47). Our current results shed a new light on the underlying adaptive process revealing that the temporal error is integrated on a trial-to-trial basis to adjust the saccade-triggering. It is noteworthy that these conclusions nicely echoe ones from saccade adaptation studies in which the adjustment of saccade amplitude has been well accounted for by postulating a Bayesian integration in which the weight associated with each piece of information is adjusted depending on the sensory evidence available (2).

Concerning the eye movements, the Kalman gain was not significantly different across the different Eye movements conditions. This denotes a consistent weight of the prediction error across different uncertainty conditions. For example, the condition of a non-stationary environment (variable speed across trials) could have encouraged a larger contribution of the final error. Note that knowledge of the magnitude of the error was not provided in the KR condition in which the speed was also interleaved. Conditions of stationary environment can be an important factor that also contributes to how the final error is weighted. In addition to stationary stimuli conditions, the temporal restrictions on which feedback is provided can also change. One limitation of our study is that we have used a relatively constant temporal window for participants to hit the moving target and the feedback was given with respect to a fixed temporal window. The learning rates or correction fractions might be also tuned to this temporal requirement and different learning rates might have been observed by varying the temporal window on which feedback was provided. For example, lax temporal constraints would lead to smaller learning rates. Recent studies have shown that different sensitivities to execution errors arise in motor learning depending on the stationarity of the environment (48). From our study we do not know whether the time course in which the two temporal errors are considered can be generalized to other conditions, such as non-stationary environments in which the temporal constraints are not constant. Future studies will have to address whether flexible learning rates also apply to the temporal domain.

## Acknowledgements

We thank Cristina de la Malla for providing feedback on an earlier version of the manuscript. This project was supported by Grants PSI2017-83493-R (AEI/Feder, UE) and 2017SGR48 from the Catalan government and PACE H2020-MSCA-ITN-2014 Ref. 642961.

## Author contribution

JLM designed the experiments. JLM, CV and LM ran the experiments and processed the data. All authors interpreted the results and wrote the article.

## Additional information

The authors declare no competing interests

